# Fatty Acyl Availability Modulates Cardiolipin Composition and Alters Mitochondrial Function in HeLa Cells

**DOI:** 10.1101/2020.02.10.937433

**Authors:** Gregor Oemer, Marie-Luise Edenhofer, Katharina Lackner, Geraldine Leman, Jakob Koch, Herbert H. Lindner, Sandrine Dubrac, Johannes Zschocke, Markus A. Keller

**Affiliations:** Institute of Human Genetics, Medical University of Innsbruck, Innsbruck, Austria; Institute of Neurophysiology, Medical University of Innsbruck, Innsbruck, Austria; Institute of Biological Chemistry, Biocenter Innsbruck, Medical University of Innsbruck, Innsbruck, Austria; Epidermal Biology Laboratory, Department of Dermatology, Venereology and Allergology, Medical University of Innsbruck, Innsbruck, Austria; Institute of Clinical Biochemistry, Medical University of Innsbruck, Innsbruck, Austria

**Keywords:** Lipids, Fatty acids, Mass spectroscopy, Cardiolipin, Mitochondria

## Abstract

The molecular assembly of cells depends not only on their balance between anabolism and catabolism, but to a large degree also on the building blocks available in the environment. For cultivated mammalian cells, this is largely determined by the composition of the growth medium used. Here we study the impact of medium lipids on mitochondrial membrane architecture and function by combining LC-MS/MS lipidomics and functional tests with lipid supplementation experiments in an otherwise serum- and lipid-free cell culture model. We demonstrate that the composition of mitochondrial cardiolipins (CL) strongly depends on the lipid environment in cultured cells and prefers the incorporation of essential linoleic acid over other fatty acids. Simultaneously, the mitochondrial respiratory complex I activity was altered, whereas the matrix-localized enzyme citrate synthase was unaffected. This suggests a link between membrane composition and respiratory capacity. In summary, we find a strong dependency of central mitochondrial features on the type of lipids contained in the growth medium. Thus, this underlines the importance of considering these factors when using and establishing cell culture models in biomedical research.

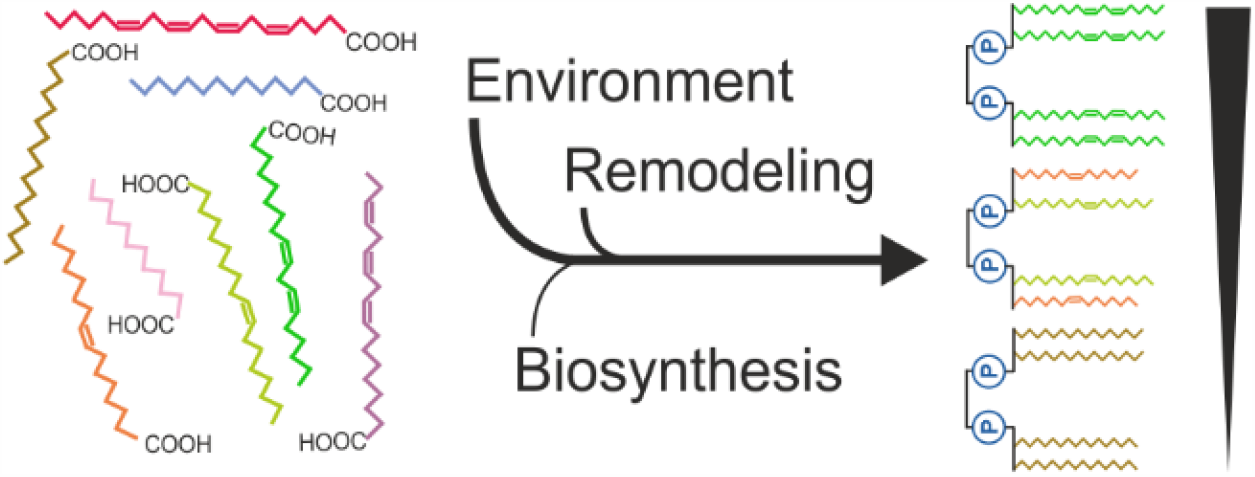

## Introduction

Immortalized human cell lines are widely used models for biochemistry, cell biology and human diseases and used in many research laboratories worldwide. They have evaded natural senescence by accumulation of proliferating mutations and can therefore be grown for prolonged periods of time *in vitro* (1). However, they require a variety of essential nutrients that is supplied with the cell culture medium. Different cell lines require different medium compositions to be viable and proliferate. For HeLa cells, the first published immortalized cell line derived from cervical cancer cells of Henrietta Lacks in 1951 (2), Harry Eagle developed his Minimal Essential Medium (MEM) (3), which was modified by Renato Dulbecco in 1959 (DMEM) (4) consisting of inorganic salts, amino acids, vitamins, D-glucose, sodium pyruvate and phenol red and is now one of the most used cell culture media (5). Additionally, also non-synthetic components are added, often derived from fetal bovine serum (FBS), which contains a complex mixture of proteins, growth factors as well as lipids and is added to the medium before usage.

To avoid batch-to-batch variation, to promote animal-free cell culture, but also to reduce costs, serum-free and chemically defined media (CDM) have been developed (6). Nevertheless, serum-containing growth conditions are still heavily used in many research laboratories. Obviously, the composition of cell culture media affects cellular composition, metabolism, proliferation, and other functions. The impact of different media components on cellular functions has been studied with variable intensity. In particular, the lipid content of the medium has received little attention in the development of recent decades. For example, most cell culture media, such as MEM, DMEM and RPMI, don’t contain any lipids and the only external source is non-chemically defined FBS, even though essential fatty acids cannot be synthesized *de novo* by human cells.

This lack of lipid-content related optimization also manifests itself in the addition of plant lipid extracts or the like in some synthetic media, which are in their fatty acid composition not necessarily matched for mimicking the original environment of mammalian cells. In previous work, we could demonstrate that HeLa cells supplemented with heart lipid extract exhibited strong lipid profile changes of mitochondrial membrane phospholipids causing a lowered basal respiration while respiratory complex I activity was increased (7). This shows that an uncontrolled composition of lipids in the cell culture medium can lead to significant functional changes. As we have recently demonstrated, in mitochondrial membranes the lipid class of cardiolipins (CLs) is especially responsive to the fatty acyl environment (8).

CLs are dimeric phospholipids with four fatty acyl side chains and are exclusively found in mitochondria. CLs have a strong preference for the inner mitochondrial membrane where they are synthesized and make up 20% of the total phospholipid content (9). The unique structure of CLs allows them to be functionally involved in mitochondrial bioenergetics (9, 10). This includes the conical shape of CL that assist with cristae structure formation, the interaction of cristae junctions with different protein complexes (11), and control of mitochondrial dynamics (12, 13). Furthermore, direct binding between CL and respiratory chain complexes is key to (super-)complex formation and stabilization (14– 16). CLs are also potent ROS scavengers, a process that leads to CL externalization to the outer mitochondrial membrane (OMM), where they recruit pro-apoptotic proteins resulting in the release of cytochrome c (cyt-c) into the cytosol and consequently apoptosis (17, 18) or mitophagy (19, 20). Importantly, the acyl side chain composition of CL is not controlled by substrate specificity of their biosynthesis, but requires an additional enzymatic remodeling processes (Figure 1A) (21, 22). In total, three enzymes have the ability to remodel CL, but only one - tafazzin - is so far reported to be pathogenic when impaired. Notably, CL compositions vary strongly across species (7), but also between tissues within one organism (8). Additionally to the observations stated above, such CL compositions have been shown to affect certain enzymatic specificities such as the cyt-c peroxidase function, which has a higher affinity towards polyunsaturated CLs (23).

**Figure 1.**
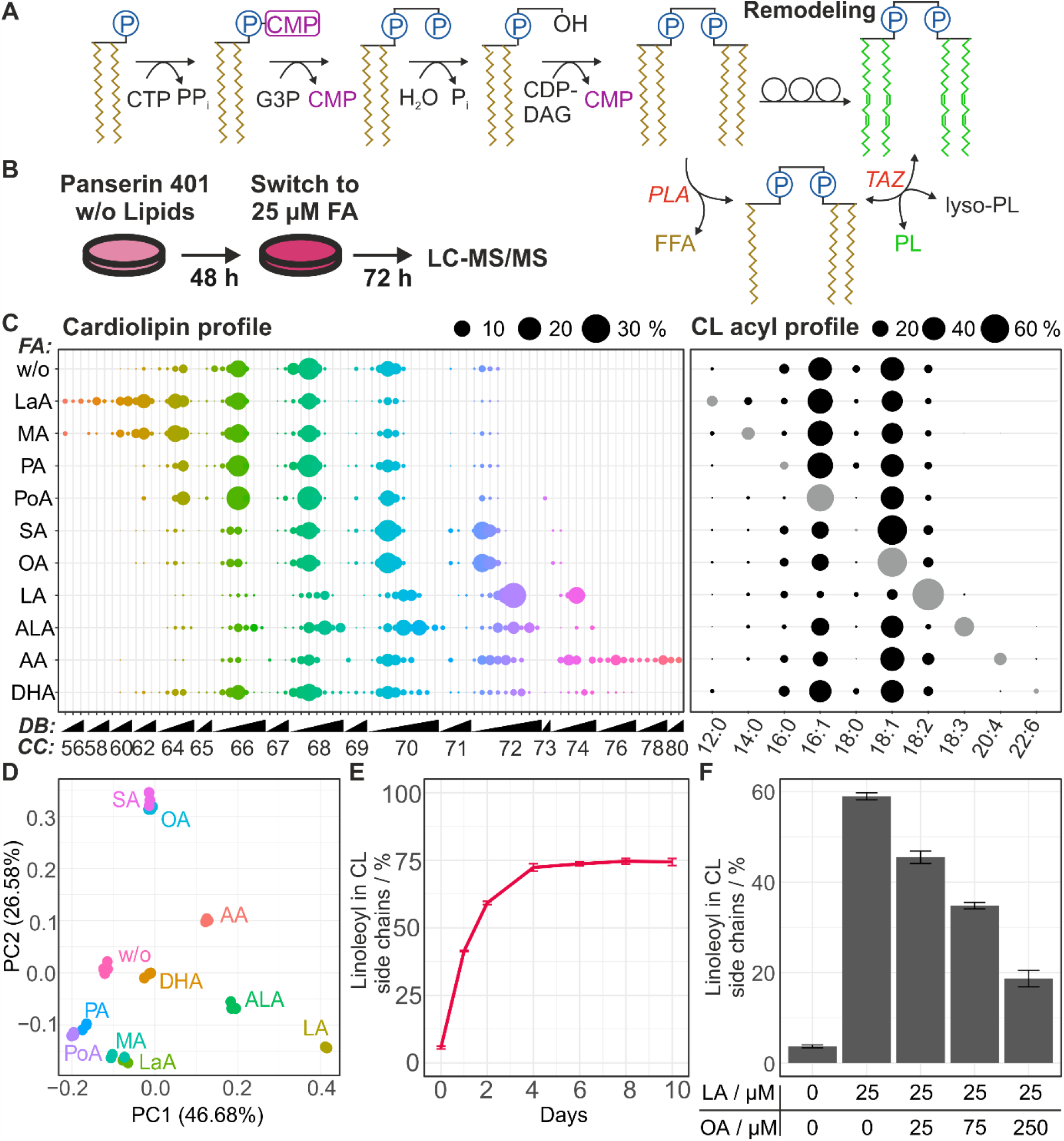
Fatty acid supplementation leads to strong fatty acyl substitution changes in CL compositions. A) CL biosynthesis and remodeling process. B) Schematic depiction of cell culture experiments. C) Mean CL profile (right) of HeLa (Mol-%) cultivated in FBS-free Panserin 401 w/o Lipids and after supplementation for 72h with 25 µM fatty acid and their respective relative CL acyl profile (left), which was obtained by utilizing MS/MS fragment spectra and mathematical modeling. DB: Total double bonds, CC: Total carbon chain length, Grey color: Fatty acid supplemented in each respective experiment. Mean ± sd profiles are shown in Figures S2 and S3. D) Principal component analysis (PCA) of CL profile showing a clear separation of supplemented conditions. Centered relative CL abundances were used for this analysis explaining over 70% of variation in the first two dimensions (PC1 and PC2). E) Linoleic acid incorporation reaches a steady-state plateau after 3-4 days in proliferating HeLa. F) Linoleic acid incorporation is clearly preferred over oleic acid, but attenuated at high excess of the latter.

Here we address the question to which extend externally added fatty acids impact on CL compositions in an otherwise lipid free environment. We investigate the speed by which the mitochondrial membrane lipid compositions adapts to changes in the fatty acyl environment and how these changes influence mitochondrial functions. To tackle this, we use a completely controlled, serum- and lipid-free cell culture model for HeLa, which can be readily supplemented with a set of fatty acids and heart lipid extract.

## Results

For establishing serum- and lipid-free cell culture, we ordered a customized Panserin 401 w/o lipids (Pan Biotech, Aidenbach, Germany) and used fatty acid supplementation mediated via fatty acid-free BSA (7). To obtain the optimal concentration for lipid supplementation, we determined the cytotoxicity of each supplemented fatty acid using a cell viability assay (CCK-8, Dojindo EU GmbH, Munich, Germany). The LD_50_-values of the 10 tested fatty acids ranged between 36.6 µM for arachidonic acid and 472.3 µM for oleic acid (see Table 1). No effect on the viability was observed with DHA in the accessible concentrations range of 0 - 200 µM which is in line with other observations (24, 25), but contrasts others (26, 27), underlining the need to determine viability for each respective model system. Due to the low LD_50_ for arachidonic acid, we used 25 µM fatty acid as a generally applicable concentration for subsequent supplementation experiments. Interestingly, these relatively low levels of supplemented fatty acids had a positive impact on cell proliferation (see Figure S1).

**Table 1.**
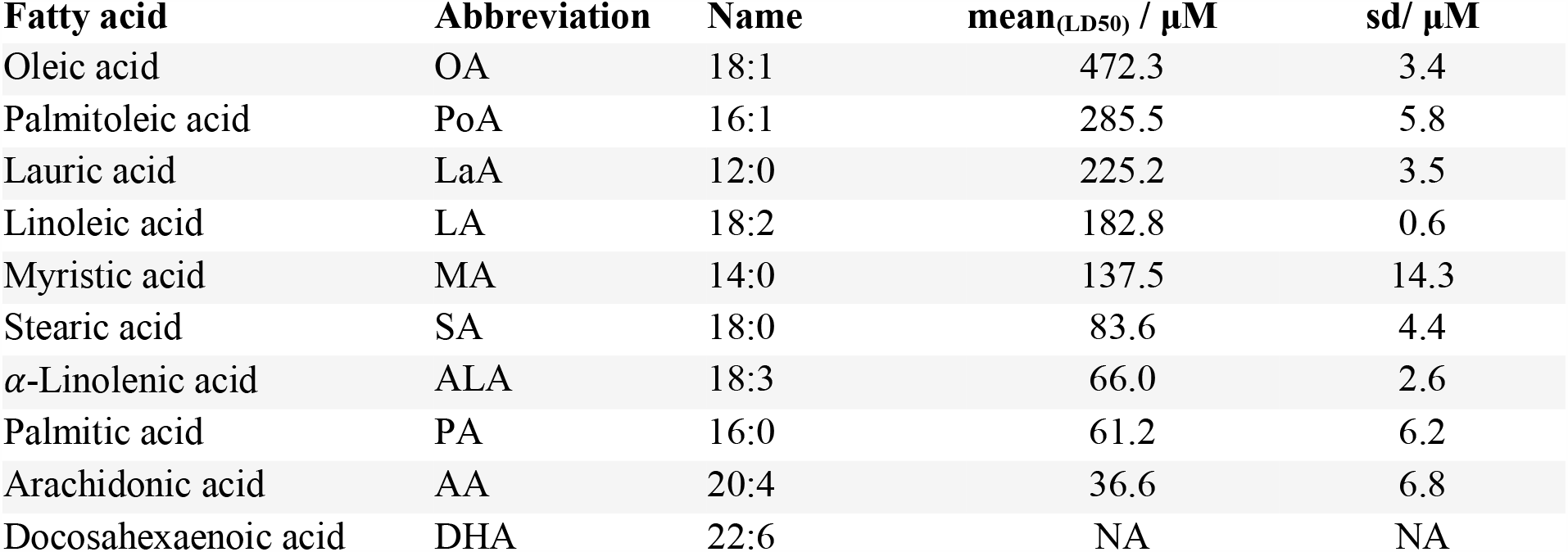
Viability of HeLa cells in the presence of supplemented fatty acid - BSA conjugates (n=3)

| Fatty acid | Abbreviation | Name | mean<sub>(LD50)</sub> / μM | sd/ μM |
| --- | --- | --- | --- | --- |
| Oleic acid | OA | 18:1 | 472.3 | 3.4 |
| Palmitoleic acid | PoA | 16:1 | 285.5 | 5.8 |
| Lauric acid | LaA | 12:0 | 225.2 | 3.5 |
| Linoleic acid | LA | 18:2 | 182.8 | 0.6 |
| Myristic acid | MA | 14:0 | 137.5 | 14.3 |
| Stearic acid | SA | 18:0 | 83.6 | 4.4 |
| α-Linolenic acid | ALA | 18:3 | 66.0 | 2.6 |
| Palmitic acid | PA | 16:0 | 61.2 | 6.2 |
| Arachidonic acid | AA | 20:4 | 36.6 | 6.8 |
| Docosahexaenoic acid | DHA | 22:6 | NA | NA |

To characterize potential lipidomic alterations caused by fatty acid supplementation, HeLa cells were grown in Panserin 401 w/o lipid for 72h in the presence of the respective fatty acid-BSA conjugate and the CL composition was quantified (Figure 1B) as described in (7). Strikingly, each cell culture medium compositions had a unique effect on the CL composition (Figure 1C). This falls in line with our earlier observation that the exact CL composition is not transcriptionally regulated but rather dependent on the availability of fatty acyls (8). In line with these reported findings, we also observe changes of the phosphatidylethanolamine and phosphatidylcholine profiles in the different conditions tested (see Figure S4).

HeLa cultivated in Panserin 401 w/o lipids can only utilize FAs derived from their own *de novo* biosynthesis and had a CL profile dominated by CL species with four double bonds (Figure 1C, left panel). Our fatty acid deconvolution revealed that palmitoleic acid (16:1, PoA) and oleic acid (18:1) made up 80% of the total CL composition, showing an apparent preference of monounsaturated long-chain fatty acids over saturated ones (Figure 1C, right panel).

A principal component analysis of the CL profiles demonstrates that the effects of supplementation are FA specific and of varying intensity (Figure 1D). The highest degree of incorporation was achieved with linoleic acid (LA) to over 70% of all side chains in favor of 16:1 and 18:1 (Figure 1C, right panel). Stearic acid (SA) supplementation led to an equally strong increase of 18:1 incorporation as treatment with oleic acid (OA) suggesting a quick desaturation process of this saturated fatty acid once taken up into the cell. Notably, palmitic acid (PA) was also not incorporated as FA16:0 itself, but led to a slight increase of FA16:1. This likely reflects the fact that palmitic acid is the major product of FA biosynthesis, and is subsequently elongated and/or desaturated into other, non-essential FA such as palmitoleic acid (PoA), stearic acid, and oleic acid (28). Myristic acid (MA), and arachidonic acid (AA) were both incorporated to a level of 11% of CL side chains and α-linolenic acid (ALA) with 27.5%. DHA supplementation led to minor changes in CL profile with levels lower than the fatty acyl resolution of the mathematical side chain modeling approach used (7). In absolute terms, upon linoleic acid treatment a total of 76% of all CL side chains were altered, followed by ALA (34.2%), AA (31.4%), and oleic acid (25.2%), while palmitic acid and DHA had the least impact with 8.9% and 8.3%, respectively (Figure S5). Notably, the amount of fatty acyl incorporation was concentration dependent. For example, when palmitic acid was supplied in excess at 200 µM - a concentration at which cell viability is already reduced to 10% within 72 h - the content of palmitic acid in CL increases from 4.6% to 40.4%, when compared to the 25 µM palmitic acid treatment (Figure S6).

Linoleic acid (18:2) was not only the most influential of the ten supplemented fatty acids, but was also causing the CL profile to focus around tetralinoleoyl-cardiolipin (TLCL), rather than leading to multiple cardiolipin compositions (see Figure 1B). The fraction of linoleic acid content in CL side chains however plateaus at 75% after four days of treatment with 25 µM (Figure 1E). Interestingly, TLCL is the most abundant molecular CL species in most mammalian tissues including heart and muscle (8). In contrast, the omega-3 α-linolenic acid (18:3) only incorporated half as frequently when being supplemented in the same concentration, highlighting a clear selectivity for linoleic acid. To check how strong the influence of a single fatty acid is in the context of dual mixtures, linoleic acid was supplemented with increasing amounts of oleic acid (Figure 1F). We observed that the amount of incorporated linoleic acid can be quenched, but a significant molar surplus of oleic acid (more than 3-fold) is needed to achieve a 50% reduction.

To test whether LA in our proliferating cell culture model remains accumulated in CLs or is readily exchanged with other FAs, once LA was removed from the medium, we performed three different lipid exchange experiments (Figure 2A). Cells were preconditioned with lipid-free (Figure 2B, upper panels), oleic acid-(Figure 2B, center panels), and α-linolenic acid-containing medium (Figure 2B, bottom panels), which was then exchanged for linoleic acid-containing medium and *vice versa*. The content of the respective fatty acyls in CL was then tracked over time by means of LC-MS/MS.

**Figure 2.**
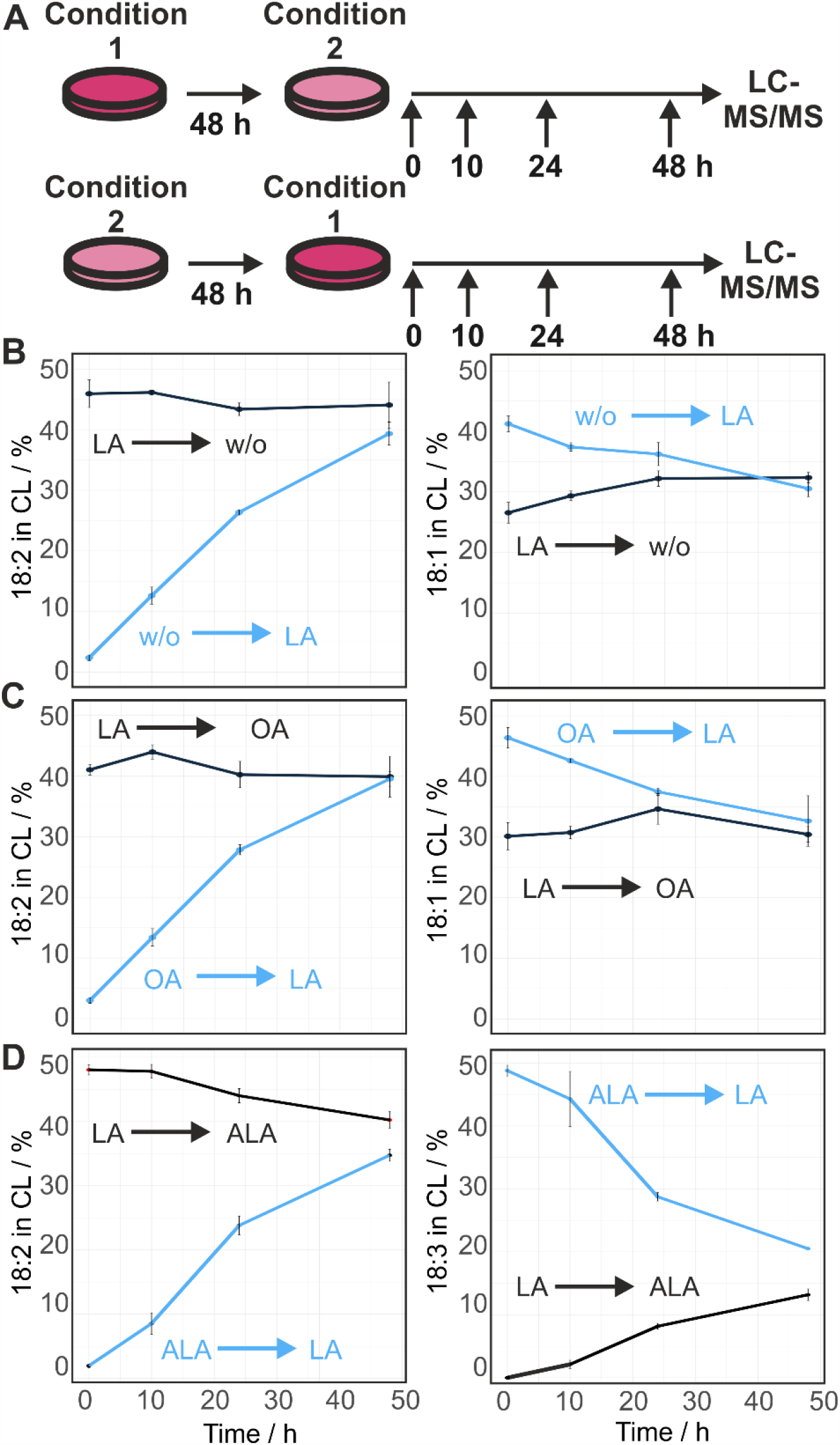
LA incorporation is preferred even if other FAs are available. A) Schematic depiction of lipid exchange cell culture experiments. B) Panserin 401 w/o and linoleic acid lipid exchange: While the amount of 18:2 increased from 2.5% to 38%, 18:1 decreased from 42% to 31% within 48h upon adding 25 µM linoleic acid. In reversed conditions, the amount of 18:2 dropped from 46% to 44% and 18:1 increased from 27% to 32%. Thus, 18:2 is relatively slowly exchanged in linoleic acid depleted conditions. C) linoleic acid (LA) and oleic acid (OA) exchange: In linoleic acid pretreated conditions, neither 18:2 nor 18:1 content was affected after replacing linoleic acid with oleic acid within 48 h. In contrast, 18:1 content decreased from 46% to 33% after oleic acid to linoleic acid replacement while LA content increased from 3% to 40% in the same time. Thus, 18:2 has a lower turnover rate in CL than 18:1 D) Linoleic acid and α-linolenic acid (ALA) exchange: Linoleic acid content decreased from 47% to 40% after ALA replacement, while alpha-linolenic acid increased from 0% to 13% in 48h. However, in reverse experiment conditions, linoleic acid increased from 3% to 34%, while alpha-linolenic acid dropped from 48% to 21%.

We observed a rapid linoleic acid incorporation in all three preconditioned scenarios, which was accompanied by a replacement of other fatty acyls including 18:1 and, when present, 18:3 (Figure 2B). Quantitatively, these effects were slightly less pronounced when exchanging linoleic acid and α-linolenic acid with each other. Upon linoleic acid withdrawal, however, the fraction of linoleyl side chains in CL remained largely stable within the 48 h time period. This effect was again least pronounced in the context of α-linolenic acid. This was surprising for this proliferating system, since at least a dilution factor of two would in theory be expected. This leads to the assumption that other cellular linoleyl reservoirs are first depleted before significant changes occur in the mitochondrial CL composition. Together this demonstrates that the CL side chain composition is highly depended on the availability of fatty acids. Thereby, the CL profile does not directly mirror the available fatty acyls, but the profile is modulated and altered by intrinsic specificities and preferences, especially for linoleic acid and to a lesser degree also for α-linolenic acid.

Next, we used this lipid- and serum-free model system to address the question how the changes in CL composition affect mitochondrial function. Previously, we demonstrated that supplementation of cultured cells with heart lipid extracts led to a more heart-like CL profile as well as an increase of respiratory complex I activity (7). Although the heart lipid extract used was composed to a considerable extent of bound and free fatty acids, and an increased proportion of this can be assumed to be linoleic acid, it could not be specified whether both effects were caused by exactly the same compound contained in the mixture. Analyzing the impact of single fatty acid-BSA treatments allowed us to determine the individual effect of changing the lipid environment on mitochondria. The enzyme activity of the inner mitochondrial membrane-localized respiratory complex I (CxI) was measured after four days of fatty acid treatment and reaching a stable CL composition. The enzymatic activity of the matrix-localized citrate synthase (CS) served as control.

We observed that the CxI activity followed a fatty acid supplement-dependent behavior. Heart lipid supplemented cells were characterized by an increased CxI activity compared to controls, while CS was unaltered, leading to an increases CxI/CS ratio (Figure 3A). When supplementing individual fatty acids, the highest specific activity was found in cells supplemented with 25 µM linoleic acid. Equally high rates were also observed for the combined treatment of 25 µM linoleic acid with 25 µM stearic acid. The latter, when supplemented alone, caused the lowest respiratory complex I activity, whereas the effect caused by the combinatorial treatment appeared to be driven by linoleic acid. Both linoleic acid containing conditions resulted in specific activities significantly higher compared to treatment with the saturated fatty acids palmitic acid and stearic acid. In contrast, no significant changes between any of the measured conditions were observed in regard to citrate synthase activity (Figure 3B), which was determined from the same cell homogenates. In summary, linoleic acid stimulated CxI activity irrespective of it being supplemented alone, or in combination. This effect is in line with earlier observations made for the treatment of HeLa cells with whole heart lipid extracts (7).

**Figure 3.**
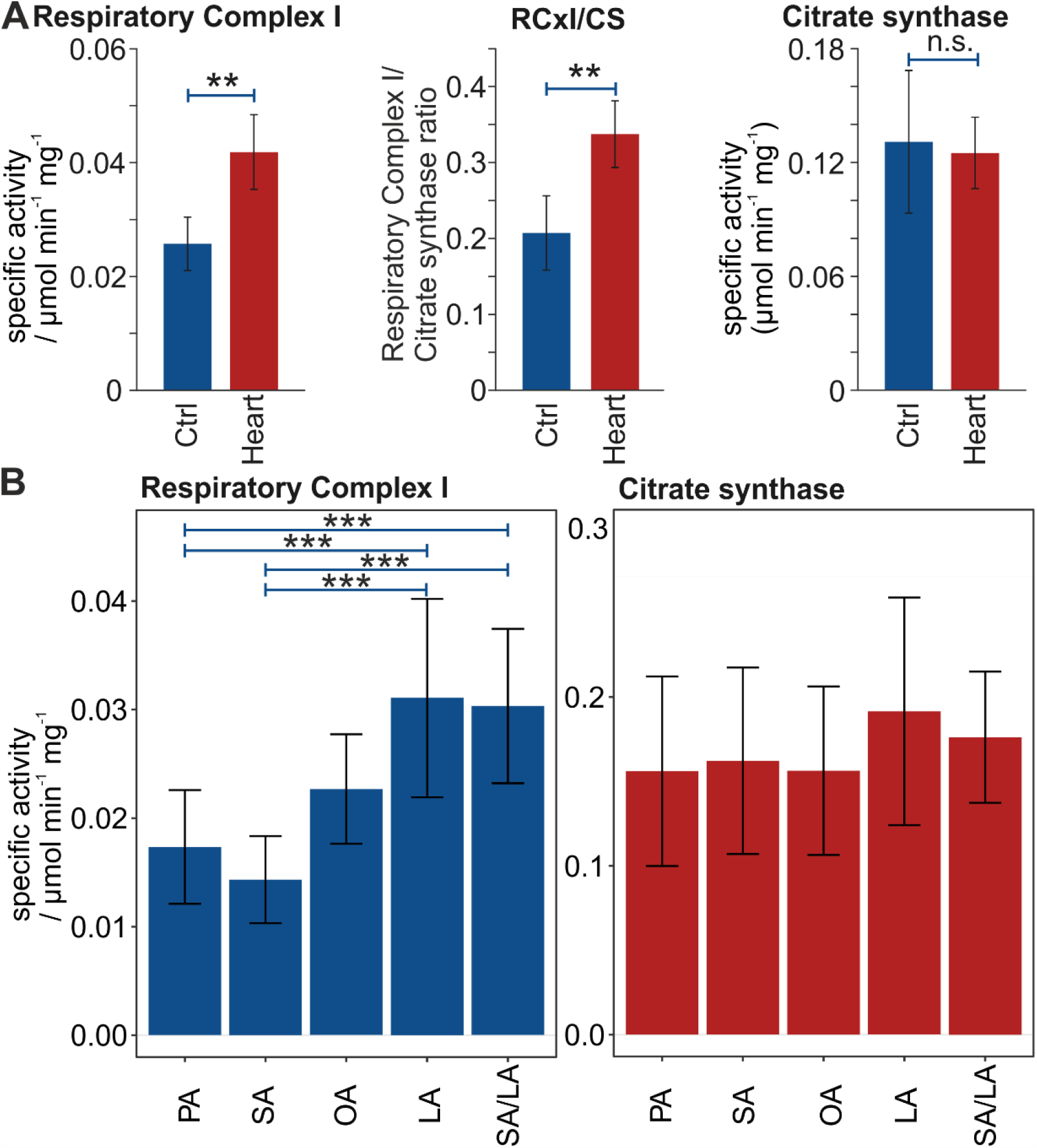
LA increases the respiratory complex I activity but not of citrate synthase. A) Respiratory Complex I (CxI) and Citrate Synthase (CS) activities in heart-lipid supplemented HeLa cells. (3 days, 200 µl/ ml heart lipid extract, n = 6, mean ± sd). B) Respiratory complex I activity was significantly increased after four days in LA supplemented media, including a 1:1 mix of LA and SA. (n = 10, 25 µM FA, mean ± sd, Significance was tested by ANOVA and Tukey post-hoc correction, (p = 7.59e-07). Citrate synthase activity was not affected by lipid supplementation in HeLa (n = 12, p = 0.454). Significance: p-value < 0.001 = ***.

## Discussion

The acyl side chain composition of CL is highly diverse in different organisms, but also within tissues of the same organism (7). The post-biosynthetic enzymatic remodeling process of CL plays an important physiological role for mitochondrial membrane constitution, which can cause severe mitochondrial dysfunction, for example when impaired as in the case of Barth Syndrome (29). It is postulated that the diversity of CL is largely driven by two processes, i) the IMM’s intrinsic strive to minimize free energy which is strongly shaped by the protein-crowded environment in the membrane (30) and ii) the set of fatty acyls available for CL remodeling that determines the local free energy minimum that can be achieved (8, 31, 32). Taken together these factors predict a strong dependence of mitochondrial membrane properties on the growth environment, which is with regard to cultured cells defined by the medium and culturing conditions chosen. Notably, we cannot exclude an effect of exogenous fatty acids on the gene expression pattern that could result in an alternative expression of respiratory complex subunits.

As demonstrated in this study, by testing the impact of a range of fatty acid supplements in cell culture, the CL compositions as well as the respiratory complex I activities respond to changes in the lipid content and composition in the medium. While the content of linoleic acid in human plasma is approximately 27% of free fatty acids, with total polyunsaturated fatty acid levels of 35-40% (33, 34), the typical lipid content of FBS mainly includes saturated and monounsaturated fatty acids with a linoleic acid fraction of as little as 5-8% (35, 36). This results in the general linoleic acid-deprived “CL phenotype” found across many different cultured cell lines (7). This does however not reflect the natural diversity of this lipid class. As a result, cells grown under these conditions tend to quickly incorporate large amounts of linoleic acid into CL once this fatty acid is made available (Figure 1E). Thereby, the degree of change that can be caused by the lipid fraction of the medium depends on the existing composition of the cells (Figure 1F and Figure 2). In this regard, a linoleic acid-saturated CL profile is noticeably more robust than other CL states.

These observations are in agreement with the finding that the turnover of CL is slower than the turnover of other phospholipids (37–39). We made the observation that the CL state in cells can be highly stable as long as the lipid environment is not altered. However, when switching from linoleic acid-depleted to linoleic-rich growth conditions we find a rapid incorporation of linoleic acid into CL (Figure 2B), reflecting the preference of CL for linolenate incorporation (40). This is reflected in the CL compositions of most tissues, especially heart, where tetralinoleyl-CL accounts for ∼80% of CL and is depleted in Barth Syndrome (29, 41).

Our observations open up a multitude of possibilities by which the medium can influence the mitochondrial CLs. These include for example batch effects caused by changes in the serum lipid composition. Also differences in the fatty acid depletion rates from the medium could lead to altered CL profiles. This would for example be the case if cells with differing growth rates are compared and linoleic acid in the medium becomes limiting for the faster growing cell line.

The same also applies to chemically defined media. Often precise information about their composition is not made public, especially with regard to the lipids used (6, 42–44). Of course, this renders it noticeably more difficult to assess to what extent the medium used is suitable for the desired cellular model system. In an ideal situation, cultivated cells could utilize the same sources and compositions of lipids as in their organism of origin. While for CL one of the main factors is the linoleic acid content, other lipid classes might be more responsive to different sets of fatty acids (8).

Another important factor is the concentration with which lipids are supplied to cultured cells. Here we chose 25 µM for all supplementation experiments as this was a concentration that generated completely viable cells, even for arachidonic acid (Table 1). Already this relatively low concentration strongly affected the side chain composition of CLs (Figure 1C). In contrast, other studies use higher concentrations ranging up to 500 µM (45, 46). Such levels can cause increased apoptosis, cell death, and the induction of endoplasmic reticulum stress (47). Not every cell culture system easily tolerates such high loads of fatty acids. Concentrations of fatty acids that impair cell viability should be avoided unless this is the exact aim of the experiment. In any case it is advisable to evaluate viability ranges not only for a given cell line in general but also in combination with its respective growth medium. Importantly, it should be noted that the observed behavior could not be transferred directly from one fatty acid to another (compare Table 1).

In a similar manner, also the solubility of fatty acids is variable, which affects the applicability for complex formation with BSA. This manifests itself by the fact that the formation of different fatty acid BSA conjugates (47) correlates with their respective melting points. Unsaturated fatty acids can be solubilized with BSA in higher concentrations than their saturated counterparts (48). Attempting to reach particularly high fatty acid concentrations could lead to increased levels of free fatty acids, which in turn can induce different physiological effects in the cell (49).

Lipids added to the cell culture medium clearly also contribute to cellular energy metabolism and anabolic requirements. We observed slightly increased growth rates of cells in the presence of low concentrations of fatty acid-BSA conjugates, irrespective of the type of fatty acid supplied (Figure S1). Such a behavior was absent in fatty acid-free BSA mock controls. It can be assumed that these effects are particularly relevant if the medium used is otherwise lipid-free, as is the case in this study. However, the variable impact of different fatty acid supplements on core mitochondrial functions as shown in Figure 3 is independent of this general behavior with regard to cell proliferation and viability.

In summary, the (lipid) composition of cell culture media strongly affects not only mitochondrial phospholipid composition but also mitochondrial functions. Supplementation, particularly with the essential linoleic acid, has a significant effect on mitochondrial activity and could be used to more realistically mimic mammalian tissue in cell culture.

## Methods

### Cell culture

If not stated otherwise, all cells were grown at 37°C, 100% humidity, and in an atmosphere of 5% CO_2_. HeLa cells were maintained in 25 cm^2^ flasks in DMEM, 1,000 mg/l D-glucose and sodium bicarbonate (D5546, Sigma Aldrich, St. Louis, USA) supplemented with 10% (v/v) heat-inactivated fetal calf serum (Fetal Calf Serum; 10500-064, Gibco, Invitrogen, Carlsbad, CA), 2 mM L-glutamine (17-605E, Lonza Group, Basel, Switzerland) and 100 U/ml Pen-Strep (17-602E, Lonza Group, Basel, Switzerland). For serum and lipid-free cell culture, medium was exchanged to customized Panserin 401 w/o lipids (Pan-Biotech, Aidenbach, Germany) for at least 2-3 days before further treatment.

### Fatty acid-BSA conjugates

For lipid supplementation fatty acid - bovine serum albumin (BSA) complexes were generated as described in (7). Briefly, fatty acids (purchased in the highest grade available from Sigma Aldrich, St. Louis, USA) were weighed and dissolved as a 100 mM stock in 100 mM NaOH. The mixture was incubated at 70°C (for stearic and palmitic acid 85°C) in a water bath until complete solubilization. Fatty acid-free BSA (A7030, Sigma Aldrich, St. Louis, USA) was dissolved in cell culture medium (2.8 mM, Panserin 401 w/o lipids), incubated at 37°C and thoroughly vortexed until complete solvation. While still warm, the 100 mM FA solution was diluted 1:5 in the 2.8 mM BSA solution (final concentration: 20 mM FA in 2.28 mM BSA) shaken for 20 min at 55°C at 1000 rpm. This BSA-complexed fatty acid stock solution was added to the cell culture medium for later supplementation (1:800 for 25 µM FA in 2.8 µM BSA solution). Medium was prepared freshly and placed at 4°C for short term storage. If not stated otherwise, the medium was exchanged every second day. PUFAs were stored at -20°C in aliquots under an argon atmosphere.

### Fatty acid supplementation of HeLa cells

For fatty acid supplementation experiments, 0.25 million cells, grown under standard conditions, were split in 6-wells with Panserin 401 w/o lipids and incubated for two days. Then, the medium was exchanged to Panserin 401 containing 25 µM of the respective supplemented fatty acid for 72 hours. Then cells were harvested by trypsinization, washed with PBS, and pelleted by centrifugation. Cell pellets were stored at -20°C for subsequent HPLC-MS/MS analysis. For fatty acid exchange experiments, 0.2 million cells were seeded in a 6-well for 48h in the Panserin 401 w/o lipids before exchanging the medium to the new target condition. Cells were pelleted as described above at the respective time points.

### Heart lipid supplementation of HeLa cells

20 g fresh pig heart (purchased at the local butchery) were homogenized in 500 ml 1x PBS using a hand blender (Bosch, Germany). The lipids were extracted following the Folch method (50) using 500 ml 2:1 chloroform/methanol mixture in a 1 l separating funnel. Organic phases containing the lipid extract were evaporated in a rotavapor at 40°C and 750 mbar. The dried lipid extracts were weighed, to 25 mg/ ml with 2:1 chloroform/methanol mixture and stored in 2 ml aliquots in glass vials at -80°C. For Heart lipid supplementation, 40 mg heart lipid extract was added to 200 ml lipid-free Panserin 401 and shaken in water bath at 50°C for 24 h (200 µg/ ml final concentration). The supplemented medium was sterile filtered through 0.2 µm pore filter. HeLa cells were grown in 25 cm2 flasks with serum-free Panserin 401 w/o lipids supplemented with pig heart lipids for three days and were then harvested.

### HPLC-MS/MS analysis of cardiolipins

Cardiolipin analysis was performed as described in (7). Briefly, sample material was homogenized in PBS and lipids were extracted following the Folch method (50) with internal standard (CL(14:0)_4_, 0.5 µM). Lipid extracts were dissolved in HPLC starting condition and subjected to HPLC-MS/MS analysis. Separation was achieved by reversed-phase HPLC with an Agilent Poroshell 120 EC-C8 2.7 µm 2.1×100mm column (Agilent Technologies, Santa Clara, USA) on a Dionex Ultimate 3000 HPLC (Thermo Fisher Scientific Inc, Waltham, USA, 50°C column oven, 0.4 µl/min flow) with running solvent A (60/40 Acetonitrile/H2O, 10 mM ammonium formate, 0.2% formic acid) and running solvent B (90/10 Isopropanol/Acetonitrile, 10 mM ammonium formate, 0.2% formic acid). Analytes were measured using a LTQ Velos MS (Thermo Fisher Scientific Inc, Waltham, USA) operated in negative ESI mode (3.8kV, 275°C capillary temperature, 460 - 1650 m/z) and data-dependent MS2 acquisition. Thermo raw data was converted to open-source MZML format and Peaks were integrated in MZmine2 (51). Identification was based on a combination of accurate mass, (relative) retention times, and fragmentation spectra, compared to a library of standards. Normalization, quantification and data analysis was performed by an in-house pipeline in R (52). CL acyl composition was modeled by a 2-step calculation: A matrix of all possible CL fragments was aligned to measured MS2 spectra by “BFGS” minimization. These CL fragments were aligned to a matrix of possible FAs and minimized using the “BGFS” algorithm. Prediction of CL profile with modeled FA and CL fragments were used as quality control. Phospholipids were measured as described in (53).

### Cell viability

Cell viability was determined using the Cell Counting Kit-8 (CCK-8, Dojindo Laboratories, Japan) in which, the reagent WST-8 [2-(2-methoxy-4-nitrophenyl)-3-(4-nitrophenyl)-5-(2,4-disulfophenyl)-2H-tetrazolium, monosodium salt] was reduced by dehydrogenases active only in viable cells to formazan, an orange soluble dye measurable by UV/Vis. Via a dilution series of each supplemented fatty acid, the LD_50_ values were calculated for each fatty acid in HeLa. Briefly, 5,000 cells suspended in 50 µl Panserin 401 were added to each well in a 96 well plate. 50 µl of same medium containing the supplemented fatty acids in different concentrations were then added and incubated for three days. Then 10 µl of the CCK-8 solution were added to each well and the plate was incubated for 2 hours at 37°C and 5% CO_2_. Finally, the absorption was measured at 450 nm by a plate reader (PHERAstar).

### Enzyme activity assays

For measuring the enzymatic activity of complex I and citrate synthase mitochondrial enriched fractions were prepared. Therefore, the cell pellet was suspended in 150 μl saccharose buffer (250 mM Saccharose, 20 mM Tris, 2 mM EDTA, 1 mg/ml BSA, pH = 7.2). After one freezing cycle in liquid nitrogen and thawing cycle at 37°C the cells were centrifuged at 12,000 rcf for 90 min. The mitochondrial cell pellet was resuspended in 150 µl hypotonic buffer (25 mmol/l KH_2_PO_4_, 5 mmol/l MgCl_2_, pH = 7.2) and further split into two aliquots: One for complex I measurement and the other for citrate synthase and protein analysis.

The citrate synthase activity was quantified by coupling the release of Coenzyme a from Acetyl-COA with the oxidation of 5,5’-dithiobis-2-nitrobenzoic acid (DTNB) and the following appearance of TNB at 412 nm in a colorimetric reaction. Briefly, a reaction mix containing 0.15 mM DTNB, 0.5 mM oxaloacetic acid and 0.1% Triton X-100 was prepared and incubated for 2 min at 30°C. For the measurement 5,000 cells were used, the reaction was started by the addition of 0.3 mM acetyl-CoA and the absorbance was recorded kinetically for 2 minutes.

The activity of respiratory complex I is directly proportional to ubiquinone reduction and the co-occurring NADH consumption was measured at 340 nm. To ensure that complex I activity was not limited by other complexes of the mitochondrial respiratory chain specific inhibitors of complex III (antimycin A) and IV (KCN) were used as well as rotenone to get the specific NADH oxidation from complex I. For complex I measurement three additional freeze/thaw cycles of the sample were performed. The reaction mix (50 mmol/l KH_2_PO_4_ pH = 7.5, 2.5 mg/ml BSA, 130 μmol/l NADH, 2 mmol/l KCN, 10 μg/ml antimycin A, 2 μg/ml rotenone) was pre-incubated for 2 min at 30°C, the reaction was started by addition of 65 µg/ml ubiquinone and the absorbance was measured kinetically for 5 min at 340 nm.

### Deposited Data

Mass spectrometric raw data and extracted lipid profiles are deposited as Mendeley Dataset (doi: 10.17632/6pdcrm5p6k.1).

## Acknowledgments

G.O. was supported by a ‘Stipendium der Monatshefte für Chemie’ of the Austrian Academy of Sciences. M.A.K. was supported by the Austrian Science Funds (FWF; P33333) and the Austrian Research Promotion Agency (FFG, #878654).

## Author contributions

G.O., J.Z., and M.A.K. designed research; G.O., M.-L.E., K.L., G.L. and M.A.K. performed research; H.L., S.D and M.A.K. contributed new reagents/analytic tools; G.O., M.-L.E., K.L., G.L., J.K. and M.A.K. analyzed data; and G.O. and M.A.K. wrote the paper with contributions from all authors.

