## Supplementary Material for "Fatty Acyl Availability Modulates Cardiolipin Composition and Alters Mitochondrial Function in HeLa Cells"

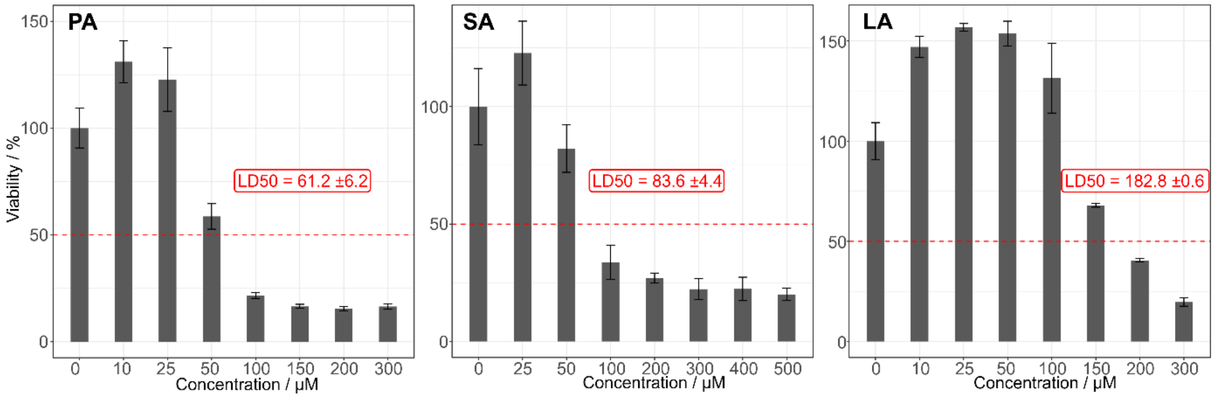


**Figure S1. HeLa cell viability and $LD_{50}$ calculations.** Each supplemented FA had a different effect on HeLa viability, yet to some extent, the viability increased at low dosage supplementation. Data is shown as mean ± sd (n= 3)

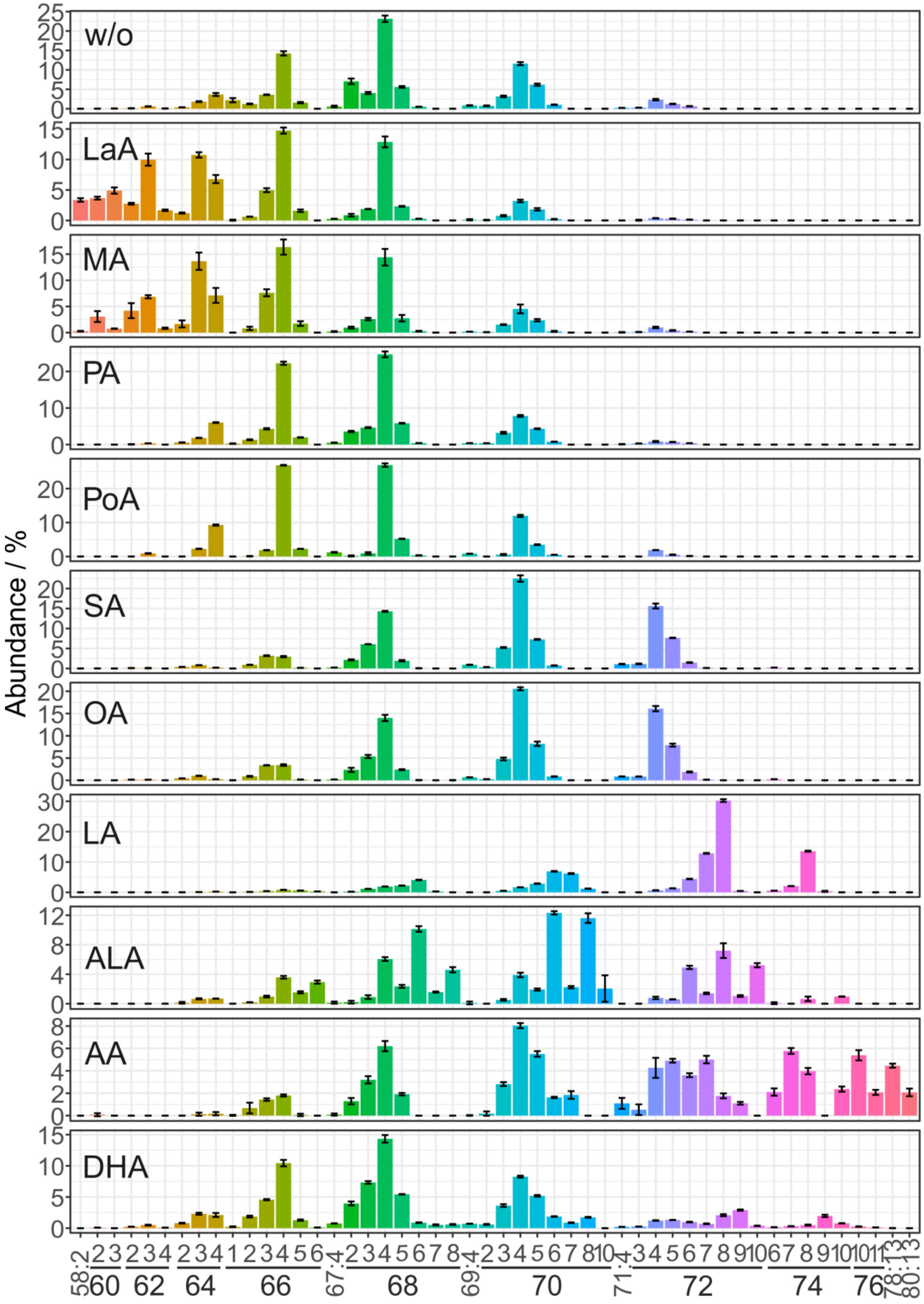


**Figure S2. Cardiolipin profiles.** Relative cardiolipin species composition in different conditions expressed as mean ± sd (biological replicates: n=3).

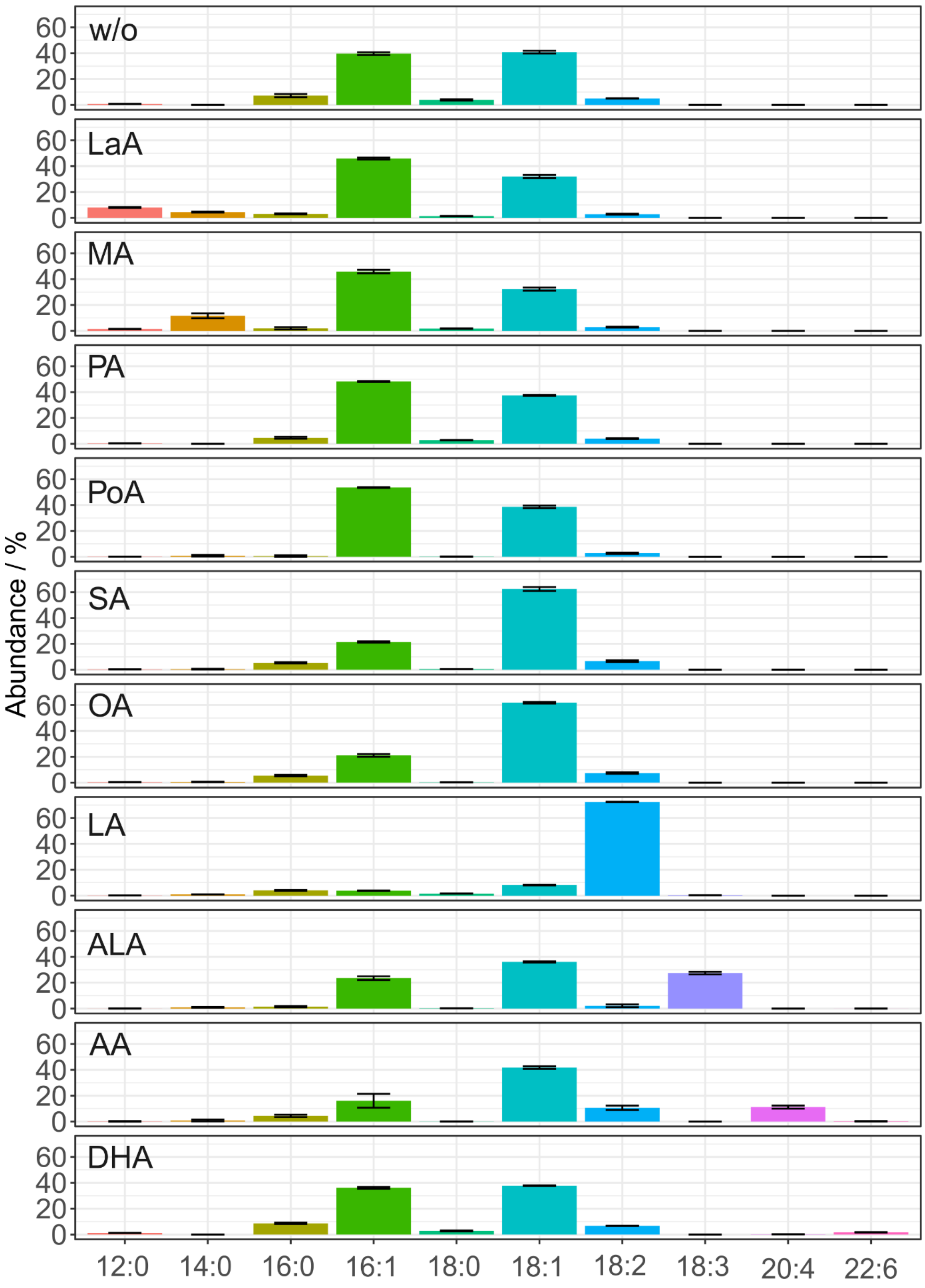


**Figure S3. Cardiolipin acyl profile modeled via MS2 spectra and FA modeling.** Relative fatty acyl substitution patterns of cardiolipin in different conditions expressed as mean ± sd (biological replicates: n=3).

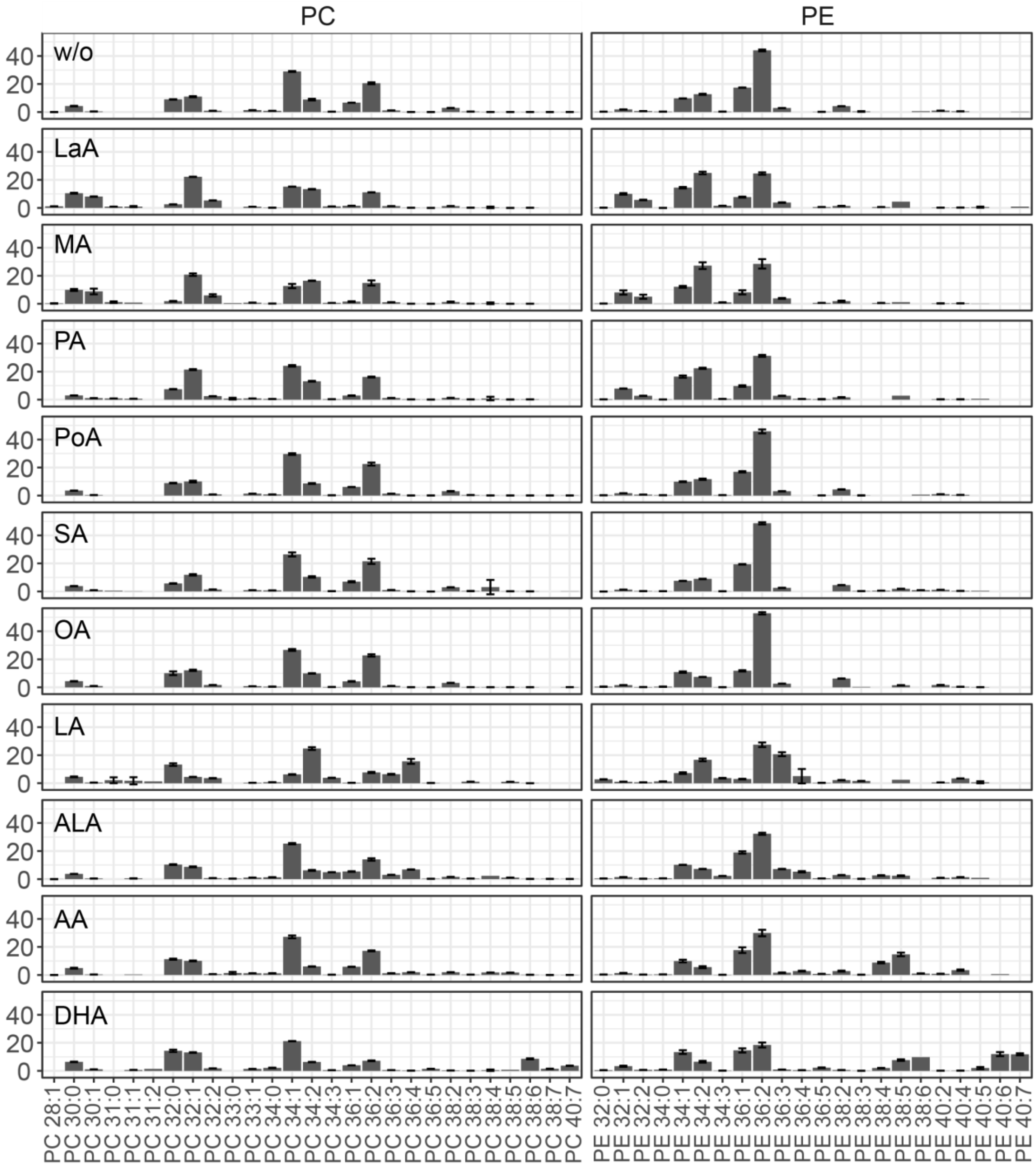


**Figure S4. Monomeric phospholipid profiles.** Left column: phosphatidylcholines (PC) and right column: phosphatidylethanolamines (PE). Class wise abundances shown in %. Lipids on the x-axis were selected for a cumulative amount of carbon atoms between 26 and 40. The shown groups distinguish between the BSA conjugated supplemented fatty acids: lauric acid (LaA), myristic acid (MA), palmitic acid (PA), palmitoleic acid (PoA), stearic acid (SA), oleic acid (OA), linoleic acid (LA), $\alpha$-linolenic acid (ALA), arachidonic acid (AA), docosahexaenoic acid (DHA). Values are shown as mean ± SD (n=3-6) of selected conditions.

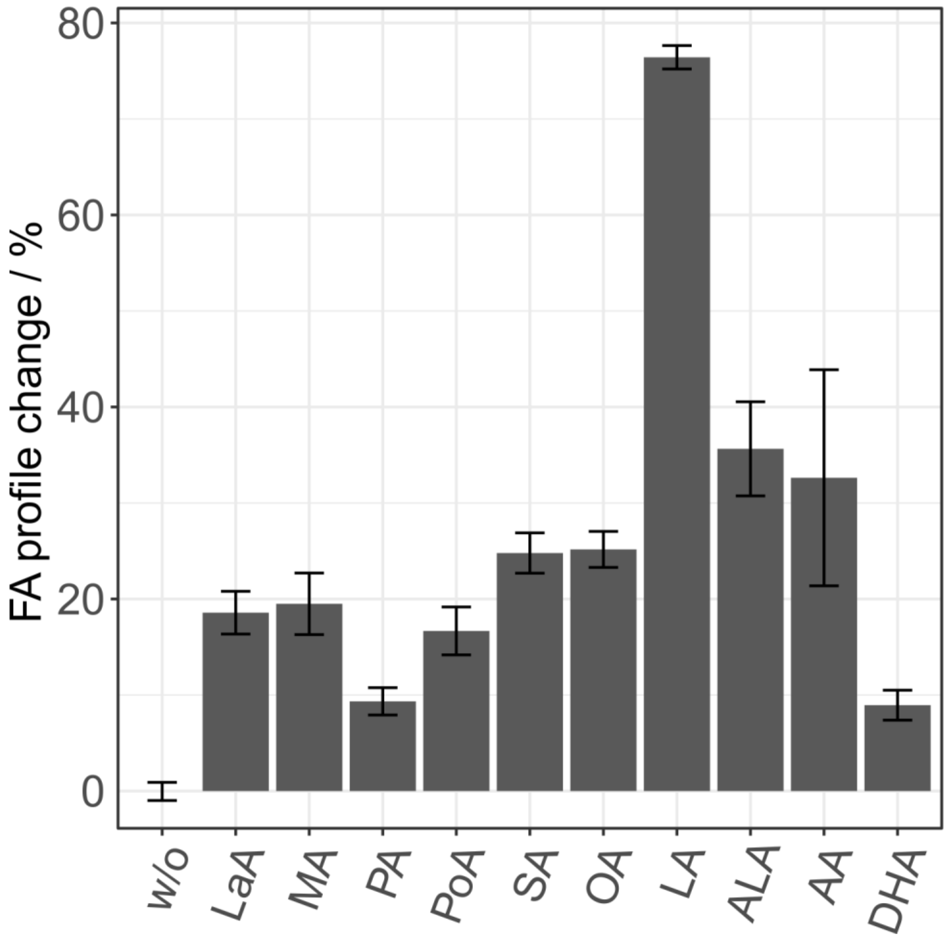


**Figure S5. Cumulative change of FA profile of CLs upon fatty acid supplementation.** Total percentage of altered cardiolipin side chains (compared to the untreated condition, w/o) in each given supplementation condition expressed as mean ± sd (biological replicates: n=3)

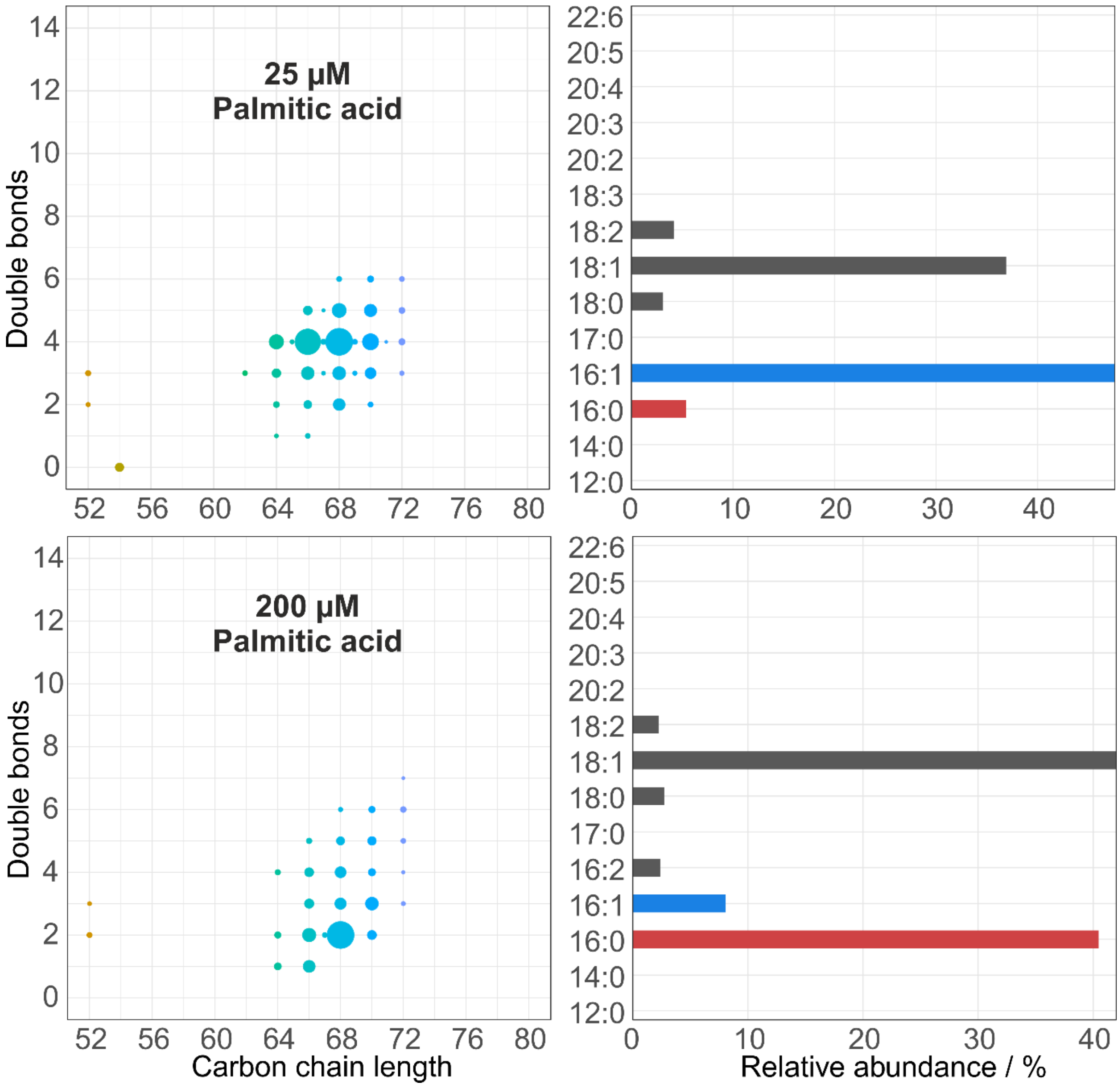


**Figure S6. CL profile and its corresponding fatty acyl side chain profile upon 25 µM (top) and 200 µM (bottom) supplementation.** Left panels: Bubble plot of cardiolipin profile in the given condition. Right panels: Relative fatty acyl substitution profiles of cardiolipins in the condition corresponding to the left panel.
